# Bacterial virulence against an oceanic bloom-forming phytoplankter is mediated by algal DMSP

**DOI:** 10.1101/321398

**Authors:** Noa Barak-Gavish, Miguel José Frada, Peter A. Lee, Giacomo R. DiTullio, Chuan Ku, Sergey Malitsky, Asaph Aharoni, Stefan J. Green, Ron Rotkopf, Elena Kartvelishvily, Uri Sheyn, Daniella Schatz, Assaf Vardi

## Abstract

*Emiliania huxleyi* is a bloom forming microalga that impacts the global sulfur cycle by producing large amounts of dimethylsulfoniopropionate (DMSP) and its volatile metabolic product dimethyl sulfide (DMS). Top-down regulation of *E. huxleyi* blooms is attributed to viruses and grazers, however, the possible involvement of algicidal bacteria in bloom demise is still elusive. We isolated from a North Atlantic *E. huxleyi* bloom a *Roseobacter* strain, *Sulfitobacter* D7, which exhibited algicidal effects against *E. huxleyi* upon co-culturing. Both the alga and the bacterium were found to co-occur during a natural *E. huxleyi* bloom, therefore establishing this host-pathogen system as an attractive, ecologically relevant model for studying alga-bacterium interaction in the oceans. During interaction, *Sulfitobacter* D7 consumed and metabolized algal DMSP to produce high amounts of methanethiol, an alternative product of DMSP catabolism. We revealed a unique strain-specific response, in which *E. huxleyi* strains that exuded higher amounts of DMSP were more susceptible to *Sulfitobacter* D7 infection. Intriguingly, exogenous application of DMSP enhanced bacterial virulence and induced susceptibility in a resistant algal strain to the bacterial pathogen. This DMSP-dependent pathogenicity was highly specific as compared to supplementation of propionate and glycerol. We propose a novel function for DMSP, in addition to its central role in mutualistic interactions, as a mediator of bacterial virulence that may regulate *E. huxleyi* blooms.

## Introduction

Phytoplankton are unicellular, photosynthetic microorganisms that contribute to about half of the estimated global net primary production, and therefore serve as the basis of the marine food web (1,2). Biotic interactions can control the fate of phytoplankton blooms in the ocean, namely predation by zooplankton, viral infections and potentially algicidal activity of bacteria (3,4). One bacterial group highly associated with phytoplankton blooms is the *Roseobacter* clade (α-Proteobacteria) (5–8) which inhabits diverse marine environments and has a wide variety of metabolic capabilities (9–11). Moreover, *Roseobacter*s were found to have a range of direct interactions, from cooperative to pathogenic, with phytoplankton species (12–16). These interactions are thought to be mediated by secreted infochemicals (17). Infochemical signaling occurs within the phycosphere, the micro-environment that surrounds algal cells where molecules can accumulate to relatively high effective concentrations (18–21). The organosulfur compound dimethylsulfoniopropionate (DMSP), and its metabolic products, plays a key role in trophic-level interactions (17) and was suggested to act as an infochemical within the phycosphere (20,22). It is produced by diverse phytoplankton species (23) and is known to mediate algae-bacteria interaction by acting as a chemoattractant (24,25) and as sulfur and carbon sources for bacterial growth (14,26,27).

*Emiliania huxleyi* is a cosmopolitan coccolithophore species which forms massive annual blooms and plays an important role in the global carbon cycle (28–31). *E. huxleyi* produces and accumulates DMSP intracellularly (up to 250 mM) (32). It harbors the gene *alma1* that encodes a DMSP-lyase responsible for high production of the volatile metabolic product dimethyl sulfide (DMS) (33). Therefore, *E. huxleyi* blooms contribute to DMS emission to the atmosphere and are thought to largely impact the global sulfur biogeochemical cycle (34,35). Once emitted to the atmosphere, DMS can undergo oxidation and induce subsequent formation of cloud condensation nuclei (36–38). The turnover of *E. huxleyi* blooms is often mediated by infection of a specific virus (*E. huxleyi* virus – EhV) that leads to rapid lysis of host cells (39–42). During the demise of *E. huxleyi* blooms an increase in bacterial abundance is observed (43–45), however, bacterial regulation of the fate of phytoplankton blooms and the cellular mechanisms governing it are largely unknown (3,8,46–48).

Activity of algicidal bacteria can be mediated by physical attachment (15,49), or by secretion of toxins or hydrolytic exo-enzymes (4,12,50) or by combing both strategies (51). For example, chemical cues from *E. huxleyi* trigger production of roseobacticides by *Phaeobacter inhibens* which leads to algal cell death (12,52). Although co-occurrence of algicidal bacteria with their algal host was demonstrated in the environment (15,49), there is still limited knowledge on how these algicidal interactions are manifested and what is their impact on phytoplankton blooms.

In the current work we isolated a *Sulfitobacter* strain (D7) from a North Atlantic *E. huxleyi* bloom. We established a robust co-culturing system in which *Sulfitobacter* D7 exhibited algicidal activity against *E. huxleyi* while consuming algal DMSP and producing high amounts of volatile organic sulfur compounds. We further examined how differential DMSP exudation, by a suite of *E. huxleyi* strains, may affect susceptibility to *Sulfitobacter* D7 infection. In a complimentary approach, we show that addition of DMSP promoted bacterial pathogenicity against *E. huxleyi* in a dose-dependent manner and induced susceptibility in a resistant algal strain. Finally, we discuss the routes in which DMSP can promote bacterial virulence and the potential role of pathogenic bacteria in regulating algal bloom dynamics.

## Results and Discussion

We obtained a bacterial consortium associated with copepods collected during an *E. huxleyi* bloom in the North Atlantic with the notion that grazers co-ingest microorganisms that interact with the algal prey (53) (Figure 1a). Inoculation of this copepod-associated microbiome (CAM) into *E. huxleyi* 379 cultures led to algal cell death. Upon application of antibiotics, the effect of CAM on *E. huxleyi* was abolished (Supplementary Figure S1a). This provided a first indication for the presence of pathogenic bacteria in CAM.

In order to study interaction of *E. huxleyi* with a specific pathogenic bacterium, we isolated from CAM a *Sulfitobacter* (termed *Sulfitobacter* D7) that belongs to the *Roseobacter* clade and sequenced its genome (BioProject PRJNA378866) (Supplementary Figures S1b and S2). *Sulfitobacter* D7 showed algicidal effects against *E. huxleyi* cultures upon co-culturing. Time course experiments of *E. huxleyi* cultures incubated with 10^3^ *Sulfitobacter* D7 mL^−1^ revealed a three-phase dynamics (Figure 1b-d). In phase I, both control and co-cultures grew exponentially, until day 9, followed by stationary phase (namely, phase II) (Figure 1b). During phase III (12–15 days) of co-culturing, algal abundance declined rapidly and algal cell death occurred in ∼90% of the population, while in control cultures it reached only ∼40% during stationary phase (Figure 1c). Rapid bacterial growth coincided with algal cell death during co-culturing, reaching 5.5·10^7^ bacteria mL^−1^ by day 16 (overall growth of four orders of magnitude), while no bacteria were observed in control cultures (Figure 1d). Interestingly, during phases II and III of co-culturing, we reproducibly detected a distinct pungent scent of volatiles that emerged only from *Sulfitobacter* D7-treated cultures (Figure 1b-d, represented by green background). *E. huxleyi* cultures incubated with CAM exhibited similar features as in co-culturing with *Sulfitobacter* D7 (Supplementary Figure S1c-e, Supplementary Text S1). Moreover, *Sulfitobacter* D7 abundance during co-culturing with CAM increased steadily by 3 orders of magnitude, as quantified by qPCR (Supplementary Figure S1e, inset). This strengthens the possible role of *Sulfitobacter* D7 as a major pathogenic component within CAM.

**Figure 1.**
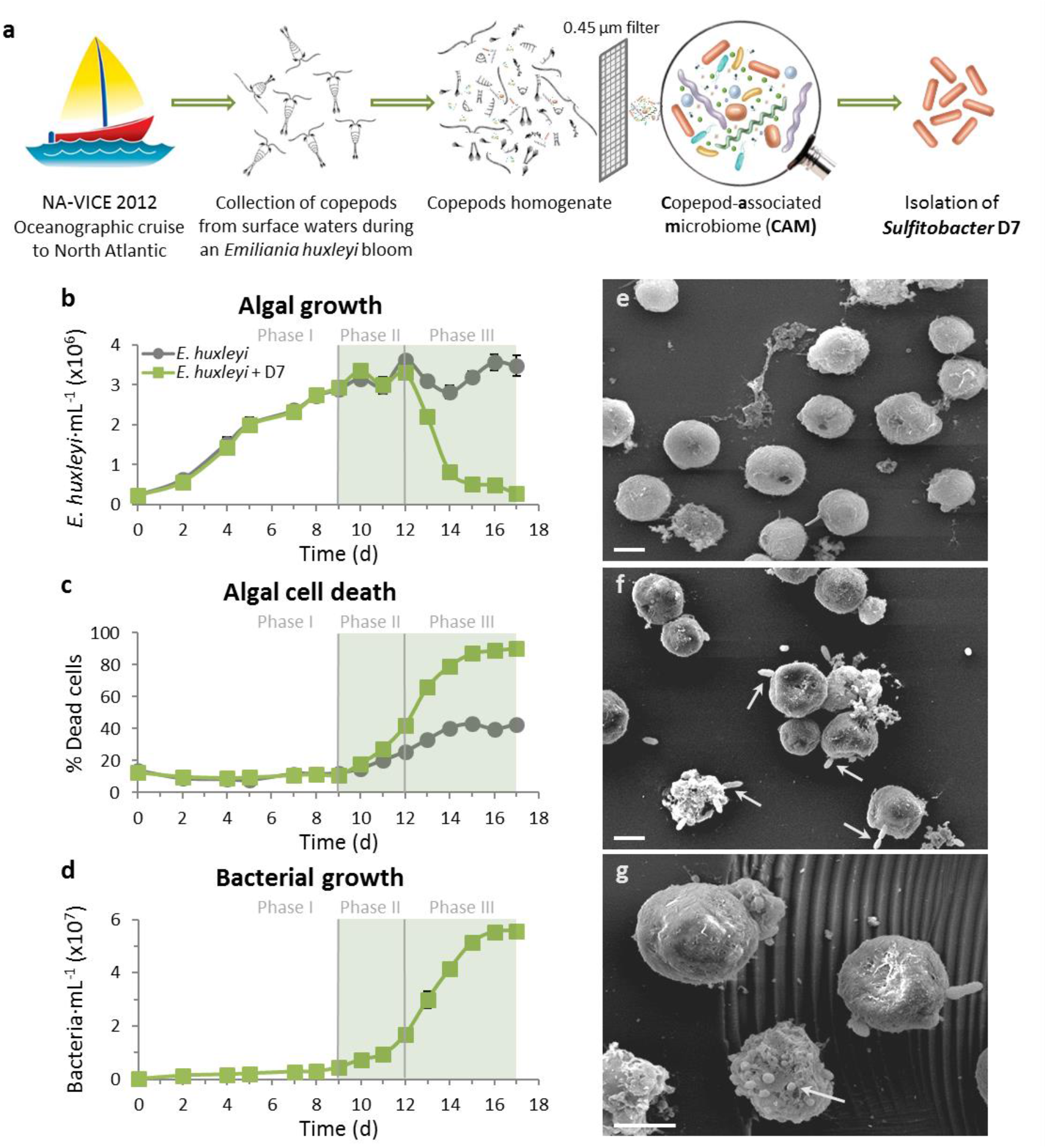
Co-culturing of *E. huxleyi* with *Sulfitobacter* D7 isolate exhibits distinct phases of pathogenicity. (a) A scheme describing the origin of the copepod-associated microbiome (CAM) bacterial consortium and isolation of *Sulfitobacter* D7 from an *E*. *huxleyi* bloom in the North Atlantic. (b-d) A detailed time course of *E. huxleyi* 379 mono-cultures (grey line) and during co-culturing with *Sulfitobacter* D7 (green line). The following parameters were assessed: (b) algal growth, (c) algal cell death and (d) bacterial growth. No bacterial growth was observed in control cultures. Green background represents the presence of a distinct scent in co-cultures. Algae-bacteria co-culturing had distinct dynamics characterized by defined phases (I-III) of pathogenicity. (e-g) Scanning electron microscopy (SEM) images of (e) uninfected *E*. *huxleyi* 379 and (f-g) *Sulfitobacter* D7-infected *E. huxleyi* cells at phase II (scale bars: 2 µm). Arrows in (f) point to *Sulfitobacter* D7 attachment to *E. huxleyi* cells. Arrow in (g) points to a membrane blebbing-like feature. Results depicted in b-d represent average ± SD (n = 3). Error bars < than symbol size are not shown. Statistical differences in (b-d) were tested using repeated measures ANOVA. *P*-values are <0.001 for the differences between control and co-cultures.

We examined the specificity of the algicidal activity of *Sulfitobacter* D7 by comparing the dynamics of co-culturing with an additional bacterial strain, *Marinobacter* D6, which was also isolated from CAM (Supplementary Figure S3). Although bacterial growth was prominent and reached similar concentrations as *Sulfitobacter* D7, the algal culture persisted in stationary growth and no increase in algal cell death was observed. Herein, we used the term “bacterial infection” to describe the algicidal impact of *Sulfitobacter* D7 on *E. huxleyi*.

Scanning Electron Microscopy (SEM) analysis of *E. huxleyi*-*Sulfitobacter* D7 interaction revealed membrane blebbing-like features in infected *E. huxleyi* cells at phase II of the infection (Figure 1g), likely corresponding to early stages of cell death (54) (Figure 1c). Furthermore, some *E. huxleyi* cells had bacteria attached to their surface in a polar manner (Figure 1f). Since previous results suggested that physical attachment of *Roseobacter*s to dinoflagellates led to algal cell death during a natural bloom (49), we speculate that *Sulfitobacter* D7 attachment to *E*. *huxleyi* cells may promote its algicidal activity, but further research on the role of physical attachment in *Sulfitobacter* D7 virulence is required.

We further aimed to explore the ecological significance of our lab-based system for alga-bacterium interaction under natural settings of a coccolithophore bloom in the North Atlantic Ocean that occurred during July 2012 (NA-VICE cruise) (42,55). We sampled the water column at different depths along a bloom patch, termed Cocco1, where we followed a water mass (quasi-lagrangian) and sampled four stations that represent a time course of ∼3 days (stations 1–4, Figure 2a, Supplementary Table S1). This allowed to locally monitor the succession of an *E*. *huxleyi* patch. Detection of *E*. *huxleyi* cells was achieved by microscopic observations, flow cytometry, molecular analyses and satellite imagery of chlorophyll fluorescence and particulate inorganic carbon (PIC), representing the calcium carbonate exoskeleton of *E. huxleyi* (42,56). *E. huxleyi* cells were prevalent in surface waters, peaking at 30–40 meters (Figure 2b-e, Supplementary Figure S4), with cell concentrations typical for oceanic *E. huxleyi* blooms (up to ∼10^3^ cells mL^−1^) (28,29). Using qPCR analyses, we detected the presence of *Sulfitobacter* bacteria which were abundant mainly at the surface, reaching a maximum level of 8.4·10^3^ bacteria mL^−1^, but was also found in deeper waters (Figure 2b-e, Supplementary Figure S4). Abundance of *Sulfitobacter* seemed to increase in surface waters along the time course while *E. huxleyi* abundance decreased between stations 1–3. The co-occurrence of *E. huxleyi* and *Sulfitobacter* in the water column during bloom succession, along with the isolation of *Sulfitobacter* D7 from the same bloom patch, demonstrates the potential existence of this algicidal interaction during *E. huxleyi* blooms. Taken together, the reproducibility of laboratory co-cultures and the natural coexistence, lay the foundation for establishing this *E. huxleyi*-*Sulfitobacter* D7 system as an attractive, ecologically relevant model for studying algicidal alga-bacterium interaction in the oceans.

**Figure 2.**
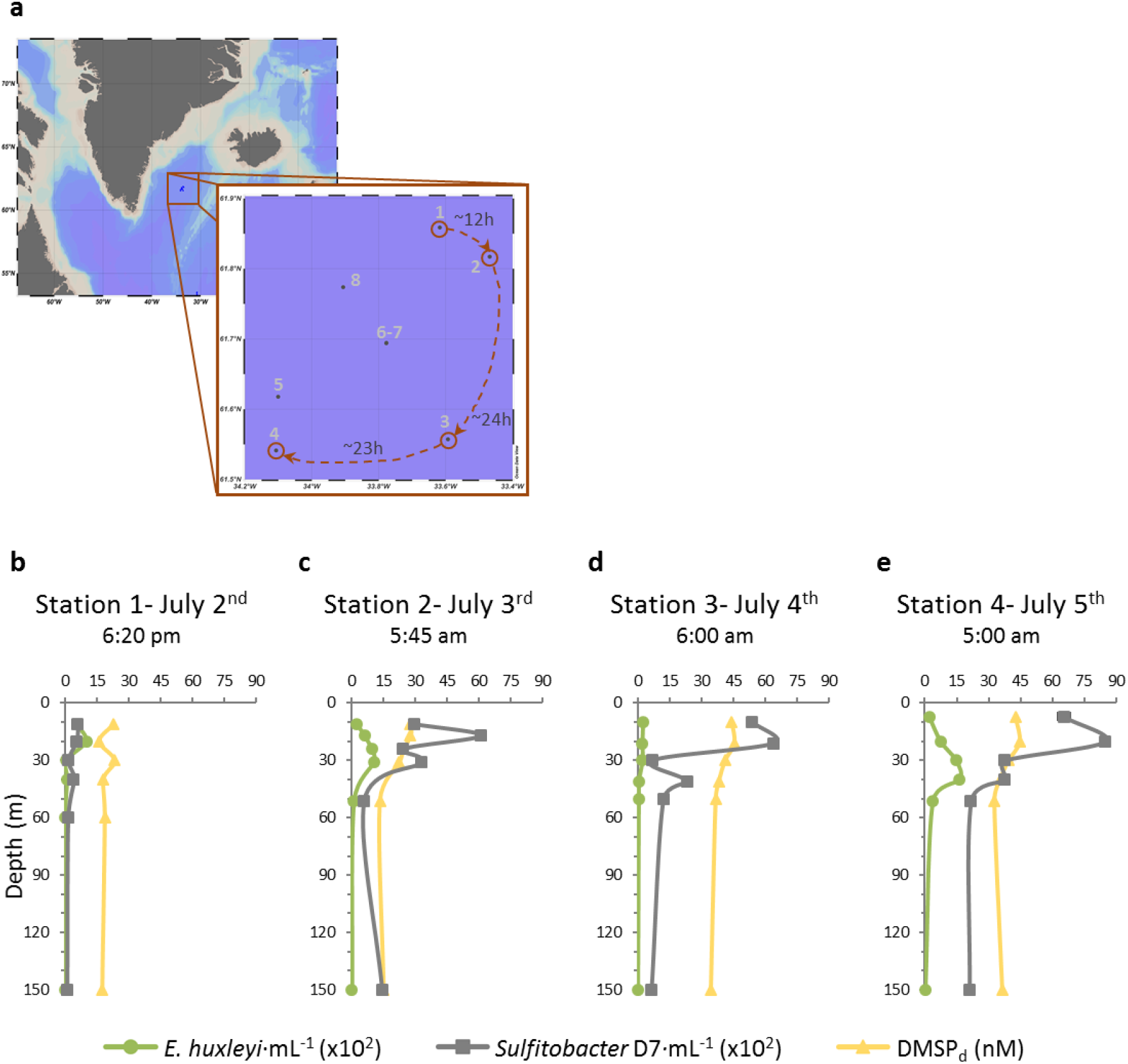
The interplay between *E. huxleyi* and *Sulfitobacter* D7 abundances and DMSPd concentration during an *E. huxleyi* bloom in the North Atlantic. (a) Sampling stations in the North Atlantic during an *E. huxleyi* bloom, July 2012 (NA-VICE cruise, 61.5–61.87°N, 33.5–34.1°W, Supplementary Table S1), in which 5–6 depths were sampled. Stations 1–4, denoted by the red track, represent a time course of the same water mass (quasi-lagrangian). Time intervals between stations are specified. (b-e) Depth profiles of *E. huxleyi* and *Sulfitobacter* D7 abundances and DMSP_d_ concentration at stations 1–4 (for stations 5–8 see Supplementary Figure S4). Results of *E. huxleyi* and *Sulfitobacter* D7 quantification represent average of three technical repeats ± SD. Error bars < than symbol size are not shown.

In order to characterize the metabolic basis of *E. huxleyi*-*Sulfitobacter* D7 interaction we sought to reveal the nature of the emitted volatiles during bacterial infection (Figure 1b-d, represented by green background). We performed an untargeted headspace-analysis using solid phase microextraction (SPME) coupled to gas chromatography mass spectrometry (GC-MS). We detected significant amounts of methanethiol (MeSH) and dimethyl disulfide (DMDS) in the headspace of *Sulfitobacter* D7- and CAM-infected *E. huxleyi* cultures, as well as small amounts of dimethyl trisulfide (DMTS) and methyl methylthiomethyl disulfide that did not appear in the headspace of control cultures (Supplementary Figure S5). A targeted analysis of the major volatile organic sulfur compounds dissolved in the media showed that DMS, MeSH and DMDS were present in *Sulfitobacter* D7-infected *E. huxleyi* cultures, while only DMS was found in control cultures (Figure 3a). The concentration of DMS in the media did not significantly differ between control and *Sulfitobacter* D7-infected cultures throughout the time course of infection (Figure 3b). In contrast, MeSH and DMDS were detected only in media of infected cultures, already in phase I, followed by a sharp increase (> 10-fold) during phases II and III (Figure 3c,d). MeSH is known to be readily oxidized to DMDS (Supplementary Figure S6c) (57) and subsequently to DMTS and methyl methylthiomethyl disulfide during sample handling (58,59). We therefore consider these volatiles part of the MeSH pool.

MeSH and DMS are known products of competing catabolic pathways of DMSP (Supplementary Figure S7) (60). The “DMSP demethylation” pathway involves enzymatic demethylation of DMSP (encoded by *dmdA* genes (61)) and subsequent production of MeSH, which can be incorporated into bacterial proteins (62). The “DMSP-cleavage” pathway is catalyzed by a DMSP-lyase enzyme (encoded by various bacterial *ddd* genes (60) and by *E. huxleyi alma1* gene (33)) and involves cleavage of DMSP and release of DMS. Since both MeSH and DMS were produced during *Sulfitobacter* D7 infection, we measured the concentration of their common precursor, dissolved DMSP (DMSP_d_), in the media of *E. huxleyi* cultures. DMSP_d_ accumulated from ∼2 µM to ∼36 µM in control *E*. *huxleyi* cultures as they aged (Figure 3e). In contrast, upon *Sulfitobacter* D7 infection DMSP_d_ concentration was comparatively low, reaching a maximal level of ∼2 µM (Figure 3e). This implies that algae-derived DMSP_d_ was consumed by *Sulfitobacter* D7 during co-culturing.

**Figure 3.**
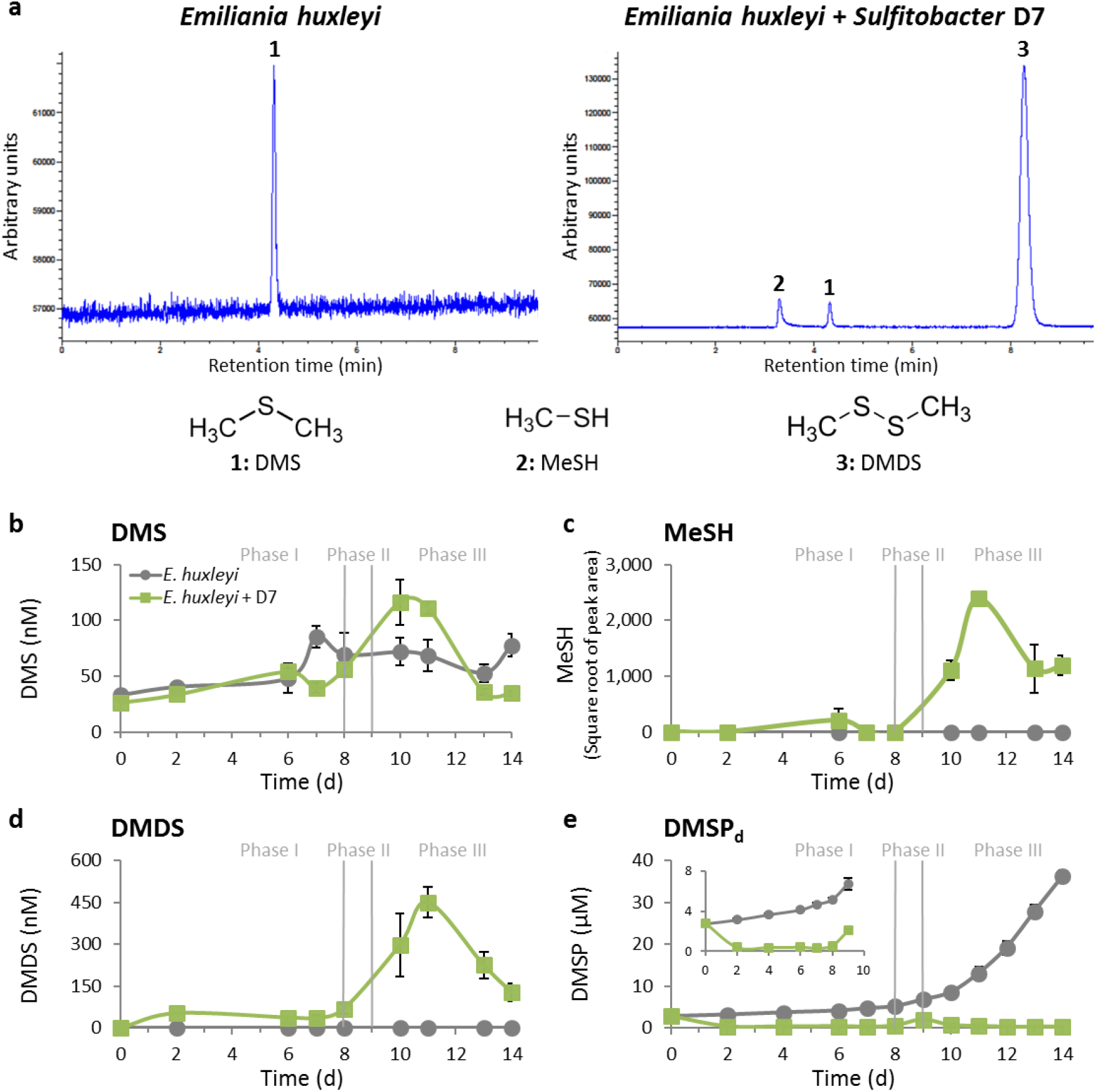
A major shift in the composition of volatile organic sulfur compounds during pathogenic phase of *Sulfitobacter* D7 infection of *E. huxleyi*. (a) Representative GC-FPD chromatograms of volatile organic sulfur compounds (VOSCs) detected in media of mono-cultures and *Sulfitobacter* D7-infected *E. huxleyi* 379 cultures at phase III (t = 11d). Peaks are marked by numbers that represent different compounds, as indicated below. DMS, dimethyl sulfide; MeSH, methanethiol; DMDS, dimethyl disulfide. DMDS is presumably an oxidation product of MeSH (Supplementary Figure S6c) and therefore considered as part of the MeSH pool. (b-e) Quantification of VOSCs; (b) DMS, (c) MeSH, (d) DMDS and (e) dissolved DMSP (DMSP_d_) in media of control (grey line) and *Sulfitobacter* D7-infected (green line) *E. huxleyi* 379 cultures during defined phases (I-III) as described in Figure 1. Inset in (e): zoom into DMSP_d_ concentration during phases I and II. Algal growth, algal cell death and bacterial growth are presented in Supplementary Figure S10. Results depicted in b-e represent average ± SD (control, n = 4; *Sulfitobacter* D7-infected, n = 2). Error bars < than symbol size are not shown. Statistical differences in (b-e) were tested using repeated measures ANOVA. *P*-values are <0.001 for the differences between control and co-cultures, except for DMS.

To identify pathways involved in DMSP catabolism, we generated a draft genome sequence for *Sulfitobacter* D7 (BioProject PRJNA378866). Gene mining analysis revealed all the putative genes of the DMSP demethylation pathway and none of the known genes in the DMSP-cleavage pathway (Supplementary Figure S7). Accordingly, *Sulfitobacter* D7 grown in mono-cultures in the presence of DMSP or in algae-derived conditioned medium consumed DMSP and produced MeSH but not DMS (Supplementary Text S2, Supplementary Table S2). We suggest that during infection *Sulfitobacter* D7 consume *E. huxleyi*-derived DMSP and produce MeSH which can be assimilated into bacterial biomass. DMS found both in control and infected *E. huxleyi* cultures was most likely a product of the activity of the DMSP-lyase, Alma1, encoded by *E. huxleyi* (33).

Interestingly, DMSP_d_ was detected during the *E. huxleyi* bloom that we sampled in the North Atlantic Ocean (Figure 2b-e, Supplementary Figure S4). The concentrations ranged between 13–45 nM, which were comparable with previous studies of *E. huxleyi* blooms (5,43). The presence of this metabolic currency along with *E. huxleyi* and *Sulfitobacter* D7 strengthens the potential of this interaction occurring in the natural environment and that it is mediated by algal DMSP.

We further aimed to assess the interplay between accumulation of algae-derived DMSP_d_ and the dynamics of *Sulfitobacter* D7 growth and pathogenicity. We used a suite of axenic *E. huxleyi* strains which differentially accumulated DMSP_d_ in media of mono-cultures (Figure 4a). This difference was most prominent in stationary phase (11 days) when media of *E*. *huxleyi* strain 379 had the highest DMSP_d_ concentration, followed by strains 1216, 373 and 2090 (72, 27, 13 and 5.5 µM on average, respectively). Inoculation of *Sulfitobacter* D7 into conditioned media (CM) derived from all *E. huxleyi* strains in stationary phase (11 days) revealed that *Sulfitobacter* D7 consumed alga-derived DMSP (Table 1) and bacterial growth was highly correlated with initial DMSP_d_ concentration (Figure 4b). CM derived from *E. huxleyi* 379 had the highest bacterial yield after 24 h of growth followed by CM from *E. huxleyi* 1216, 373 and lastly 2090 (1.1·10^8^, 7.5·10^7^, 2·10^7^ and 1.7·10^7^ bacteria mL^−1^ on average, respectively) (Figure 4b). Intriguingly, we detected substantial variability in infection dynamics among *E. huxleyi* strains (Figure 4c-f). All strains were infected at various degrees, presenting all three phases of pathogenicity (Figure 4c-e), except for *E*. *huxleyi* 2090 that was unaffected by the presence of bacteria (Figure 4f, Supplementary Table S3). Pronounced differences were observed for the dynamics of phase III in which *E. huxleyi* 379 cultures declined most rapidly (within 5 days) followed by strains 1216 (within 7 days) and 373 (within 10 days). In all cases, the decline in algal abundance correlated with growth of bacteria that reached ∼10^8^ bacteria mL^−1^, corresponding to phase III, except in 2090 where bacterial abundance was 10-fold lower. Moreover, the duration of phase III had an inverse correlation with the concentration of DMSP_d_ in the media of uninfected *E. huxleyi* strains (Supplementary Table S3). Namely, strains that accumulated more DMSP_d_ in the medium during algal growth in mono-cultures were more susceptible to *Sulfitobacter* D7 infection during co-culturing. This raised the hypothesis that DMSP is not only an important carbon and reduced sulfur source for bacterial growth but may also promote *Sulfitobacter* D7 pathogenicity against *E. huxleyi*.

**Table 1.** Concentration of dissolved DMSP (DMSPd) and its consumption by *Sulfitobacter* D7 grown for 24 h in conditioned media (CM) derived from various *E. huxleyi* strains at 11 days of growth (Figure 4a).

|  | DMSP<sub>d</sub> (μM) |  | DMSP consumed (μM)<sup>d</sup> |
| --- | --- | --- | --- |
|  | T=0h<sup>a</sup> | T=24h<sup>b</sup> |  |
| CM-379 | 71.3 | 44.4 ± 1.6 | 26.9 |
| CM-1216 | 26.5 | < 0.15<sup>c</sup> | 26.3<sup>e</sup> |
| CM-373 | 13 | < 0.15<sup>c</sup> | 12.8<sup>e</sup> |
| CM-2090 | 5 | 3.4 ± 0.01 | 1.6 |

<sup>a,b</sup> Results represent average ± SD (<sup>a</sup>n = 1, <sup>b</sup>n = 3)
<sup>c</sup> Not detected. Detection limit was 150 nM
<sup>d</sup> Estimated by subtraction of the concentration at 24 h from 0 h
<sup>e</sup> Under estimation. Due to detection limits we assumed concentration of 150 nM at T=24h

**Figure 4.**
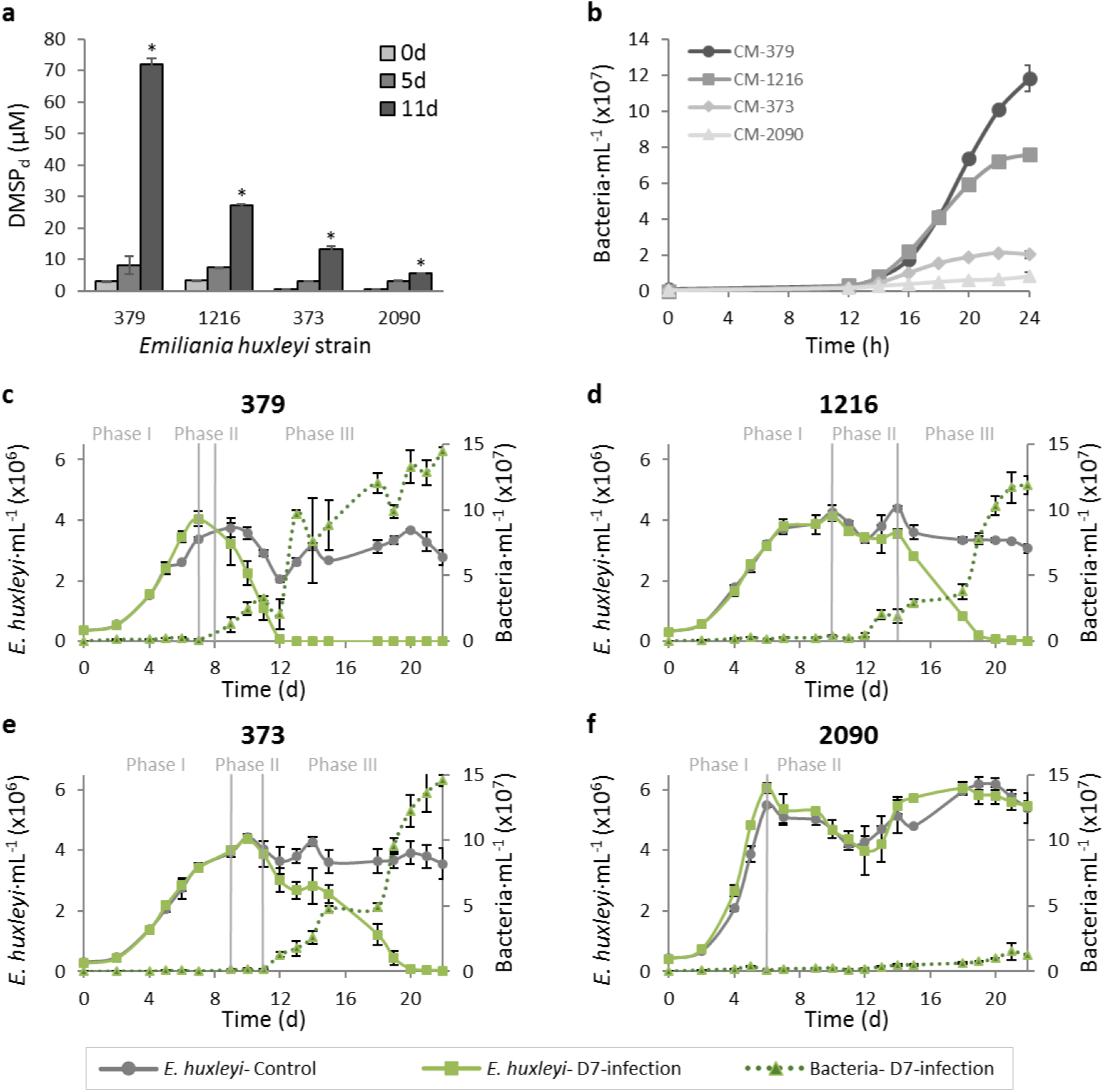
*E*. *huxleyi* strain-specific DMSP exudation and susceptibility to *Sulfitobacter* D7. (a) Concentration of dissolved DMSP (DMSP_d_) in media of mono-cultures of four axenic *E. huxleyi* strains (379, 1216, 373 and 2090) at different stages of growth. (b) Growth curves of *Sulfitobacter* D7 in conditioned media (CM) obtained from *E. huxleyi* cultures from panel (a) at 11 days of growth. (c-f) Differential dynamics of co-cultures of *Sulfitobacter* D7 with a suite of *E*. *huxleyi* strains. Time course of algal and bacterial growth (left and right axes, smooth and dashed lines, respectively) in mono-cultures (gray) and *Sulfitobacter* D7-infected (green) cultures of *E. huxleyi* strains (c) 379, (d) 1216, (e) 373 and (f) 2090. No bacterial growth was observed in control cultures. Defined phases (I-III) of pathogenicity are denoted. Results represent average ± SD (n = 3). Error bars < than symbol size are not shown. Statistical differences between strains in (a) were tested using one-way ANOVA for each time point, followed by a Tukey post-hoc test. * *P*-values in (a) are <0.001 for the differences between all strains on day 11. *P*-values in (b-f) were calculated using repeated measures ANOVA, followed by a Tukey post-hoc test. *P*-values in (b) are <0.001 for the differences between all CM. *P*-values in (c-e) are <0.001 for the differences between control and co-cultures. *P*-value in (f) is <0.001 only for the differences in bacterial growth between control and co-cultures.

Intriguingly, addition of DMSP to *E. huxleyi* 379 cultures inoculated with *Sulfitobacter* D7 expedited the dynamics of infection in a dose dependent manner (Supplementary Figure S8). Co-cultures supplemented with 500 µM and 100 µM DMSP collapsed after 5 and 7 days, respectively, while co-cultures in which DMSP was not added declined only after day 11. Algal mono-cultures were not affected by the addition of DMSP. In order to test the specificity of DMSP in promoting bacterial virulence we supplemented algal mono- and co-cultures with additional 3-carbon substrates, glycerol and propionate (Figure 5). Once again addition of DMSP promoted *Sulfitobacter* D7 infection dynamics, while glycerol and propionate had a minor effect (Figure 5a). The DMSP-supplemented co-cultures reached phase II after only 4 days and completely collapsed at day 8. The co-cultures supplemented with glycerol and propionate had similar dynamics as co-cultures with no substrates addition, all entered phase II at day 5 and fully collapsed at day 12. Interestingly, bacterial growth in all co-cultures were similar until day 5, with slightly more bacteria in the glycerol-treated co-cultures at day 3 (Figure 5b). Although bacterial density was similar between all the substrate-supplemented cultures the early virulence of *Sulfitobacter* D7 was invoked only in the presence of DMSP. These results provide a direct link between DMSP and algicidal activity of *Sulfitobacter* D7 against *E. huxleyi*.

We further hypothesized that the observed resistance of *E. huxleyi* 2090 to *Sulfitobacter* D7 infection may be explained by the low level of DMSP_d_ in the media of 2090 cultures (Figure 4a). Therefore, we added exogenous DMSP to *E. huxleyi* 2090 and examined its susceptibility to bacterial infection. Intriguingly, algal growth arrest was induced at day 4 of *E. huxleyi* 2090-*Sulfitobacter* D7 co-cultures supplemented with 100 µM DMSP (Figure 5c). Glycerol and propionate did not affect the dynamics of co-cultures at all, although bacterial growth was more prominent compared to the non-supplemented co-cultures (Figure 5d). High inoculum of *Sulfitobacter* D7 did not affect *E. huxleyi* 2090 growth, unless DMSP was present (Figure 5c). This strengthens the pivotal role of DMSP in mediating *Sulfitobacter* D7 virulence towards *E. huxleyi*.

**Figure 5.**
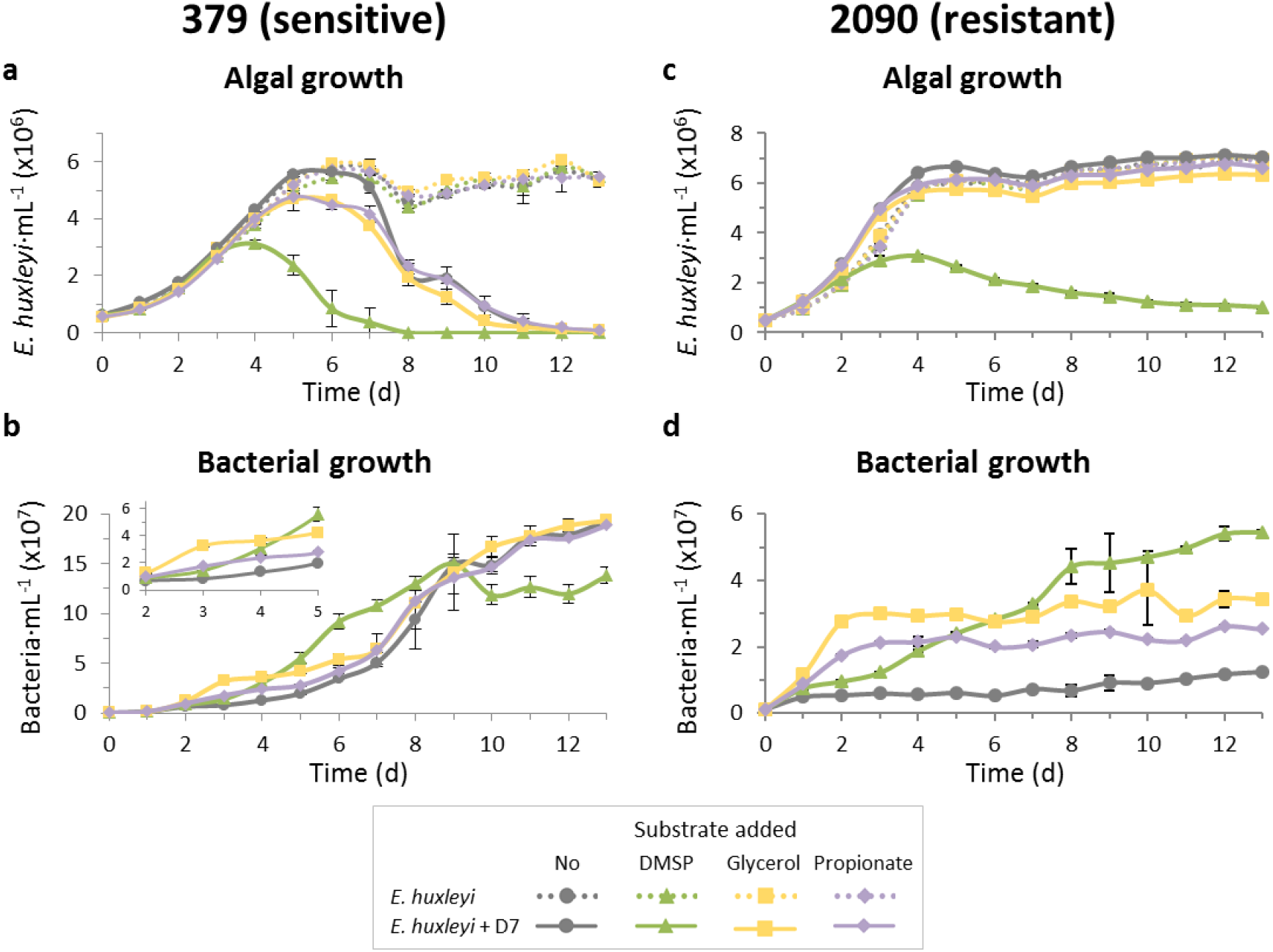
DMSP promotes *Sulfitobacter* D7 virulence towards *E. huxleyi*. Time course of algal and bacterial growth in cultures of (a-b) the sensitive *E. huxleyi* strain 379 and (c-d) the resistant *E. huxleyi* strain 2090, mono-cultures (dashed lines) and during co-culturing with *Sulfitobacter* D7 (smooth lines). Cultures were supplemented at day 0 with 100 µM of the following substrates: DMSP (green, triangle), glycerol (yellow, square), propionate (purple, diamond) or none (gray, circle). Inset in (b): zoom into bacterial growth at days 2 to 5. No bacterial growth was observed in control cultures. Results represent average ± SD (n = 3). Error bars < than symbol size are not shown. Statistical differences were tested using two-way repeated measures ANOVA, accounting for infection and the different substrates. *P*-values in (a) are <0.001 for the differences between control and co-cultures and for the differences between the DMSP treatment and the other treatments in co-cultures. *P*-values in (b) are <0.05 only for the differences between the glycerol treatment and the rest of the treatments in co-cultures. *P*-values in (c) are <0.001 for the differences between the DMSP treatment in co-cultures and the other treatments. *P*-values in (d) are <0.01 for the differences between all treatments in co-cultures.

DMSP has many suggested cellular functions including osmoregulation, antioxidant and chemoattractant (24,63,64). Our results place DMSP as mediator of bacterial virulence via several suggested cellular pathways (Figure 6). Firstly, DMSP promotes growth of *Sulfitobacter* D7 (Supplementary Figure S9) and it is consumed and metabolized to MeSH by the bacterial demethylation pathway (Figure 3). DMSP-degrading bacteria can produce more cellular energy from demethylation rather than cleavage of DMSP (65). In both pathways there is gain of reduced carbon but only through demethylation there is gain of reduced sulfur that can be incorporated into amino acids (methionine and cysteine) (62,66). As demonstrated in *P. inhibens*, these amino acids can subsequently be incorporated into bacterial algicides, roseobacticides, that kill *E. huxleyi* cells (12,52,67). Therefore, DMSP and its metabolic products can promote bacterial virulence by acting as precursors for the synthesis of bacterial algicides.

Interestingly, roseobacticides biosynthesis by *P. inhibens* is regulated by quorum sensing (QS) (52), as is virulence of many other pathogenic bacteria (68). Moreover, the production of QS molecules in *Roseobacter*s can be stimulated by DMSP (22). Thus, the involvement of QS may also be applicable in the *E. huxleyi*-*Sulfitobacter* D7 system described here. Genomes of *Sulfitobacter* spp., including *Sulfitobacter* D7, indeed encode genes involved in *N*-acyl-l-homoserine lactone (AHL)-based QS (69,70). It was also shown that a precursor for QS molecules produced by the bacterium *Pseudoalteromonas piscicida* induced mortality of *E. huxleyi* in cultures (71). Therefore, biosynthesis of QS molecules can regulate expression of virulence-related genes and may also contribute to pathogenicity by producing intermediate compounds that function as algicides themselves. Further investigation is needed in order to assess the involvement of QS and algicides in the pathogenicity of *Sulfitobacter* D7.

DMSP can also mediate *Sulfitobacter* D7 virulence by acting as a chemotaxis cue towards *E. huxleyi* phycosphere. Marine bacteria can sense DMSP and use it as a signal for chemotaxis (24) in pathogenesis (72) and symbiosis (25,73). *Sulfitobacter* D7 is a motile bacterium and its genome encodes flagella biosynthesis genes. Therefore, DMSP released from *E. huxleyi* cells can serve as a cue in which *Sulfitobacter* D7 can locate algal cells and subsequently attach and consume DMSP (Figure 1f,g, Figure 3e). Since DMSP is not a specific metabolite for *E. huxleyi*, and can be produced by diverse algal species, it is likely that other infochemicals also mediate the specificity of this interaction. Such infochemical could convey information regarding the physiological state of the algal cell. For example, *p*-cumaric acid, a molecule released from senescing *E. huxleyi* cells and was shown to trigger production of roseobacticides by *P. inhibens* (12,52).

Specificity in DMSP signaling can also be achieved by differential exudation rates among *E*. *huxleyi* strains (Figure 4a). Indeed, we found correlation between patterns of DMSP exudation and the response of *E. huxleyi* strains to *Sulfitobacter* D7 infection. Strains exhibiting higher exudation were more susceptible and died faster upon *Sulfitobacter* D7 infection (Figure 4). Therefore, the extent of metabolites exudation by algal strains would shape an “individual phycosphere”, which can potentially determine the susceptibility to bacterial infection. This algicidal microscale interaction may shape the population of *E. huxleyi* strains during algal bloom dynamics.

*E. huxleyi* bloom demise is thought to be mediated by viral infection (39–42). It is therefore raising the intriguing question of how come *E. huxleyi* strains resistant to viral infection do not take over the bloom under viral pressure. Our study reveals an important algicidal control by bacteria that possibly constrain the outgrowth of virus-resistant *E*. *huxleyi* strains. Strains of *E. huxleyi* resistant to viral infection (373, 379) (74) were highly susceptible to *Sulfitobacter* D7. Conversely, *E*. *huxleyi* 2090, which is highly susceptible to viral infection (74), was resistant to *Sulfitobacter* D7. We propose that a tradeoff between susceptibility to viral infection and bacterial pathogenicity, mediated by DMSP, may affect the fate of *E. huxleyi* cells during bloom dynamics. Moreover, lysis of *E. huxleyi* cells by viral infection leads to release of dissolved organic matter, including DMSP (75), which in turn can boost bacterial growth and virulence of pathogens, such as *Sulfitobacter* D7 (5,76). Therefore, algae-bacteria interaction may have an underappreciated active role in phytoplankton bloom demise. Further research is required in order to assess the impact of algicidal bacteria on phytoplankton bloom dynamics. Determination of algal bloom demise dominated by viruses or bacteria would encompass many challenges. Mechanistic understanding of these microbial interactions would be essential in order to assess their relative metabolic and biogeochemical imprint.

*E*. *huxleyi* blooms are an important source of DMS emission (34,77). The balance between competing DMSP catabolic pathways, driven by microbial interactions (bacterium-bacterium, alga-bacterium, alga-virus), may regulate oceanic sulfur cycling (Figure 6) (75,78). Interactions of algae with pathogenic bacteria may shunt DMSP catabolism towards high amounts of MeSH, on the expense of DMS, and can boost bacterial growth by incorporation of this reduced sulfur and carbon source. This metabolic switch may constitute a profound biogeochemical signature during algal blooms by affecting the cycling of sulfur and feedback to the atmosphere.

**Figure 6.**
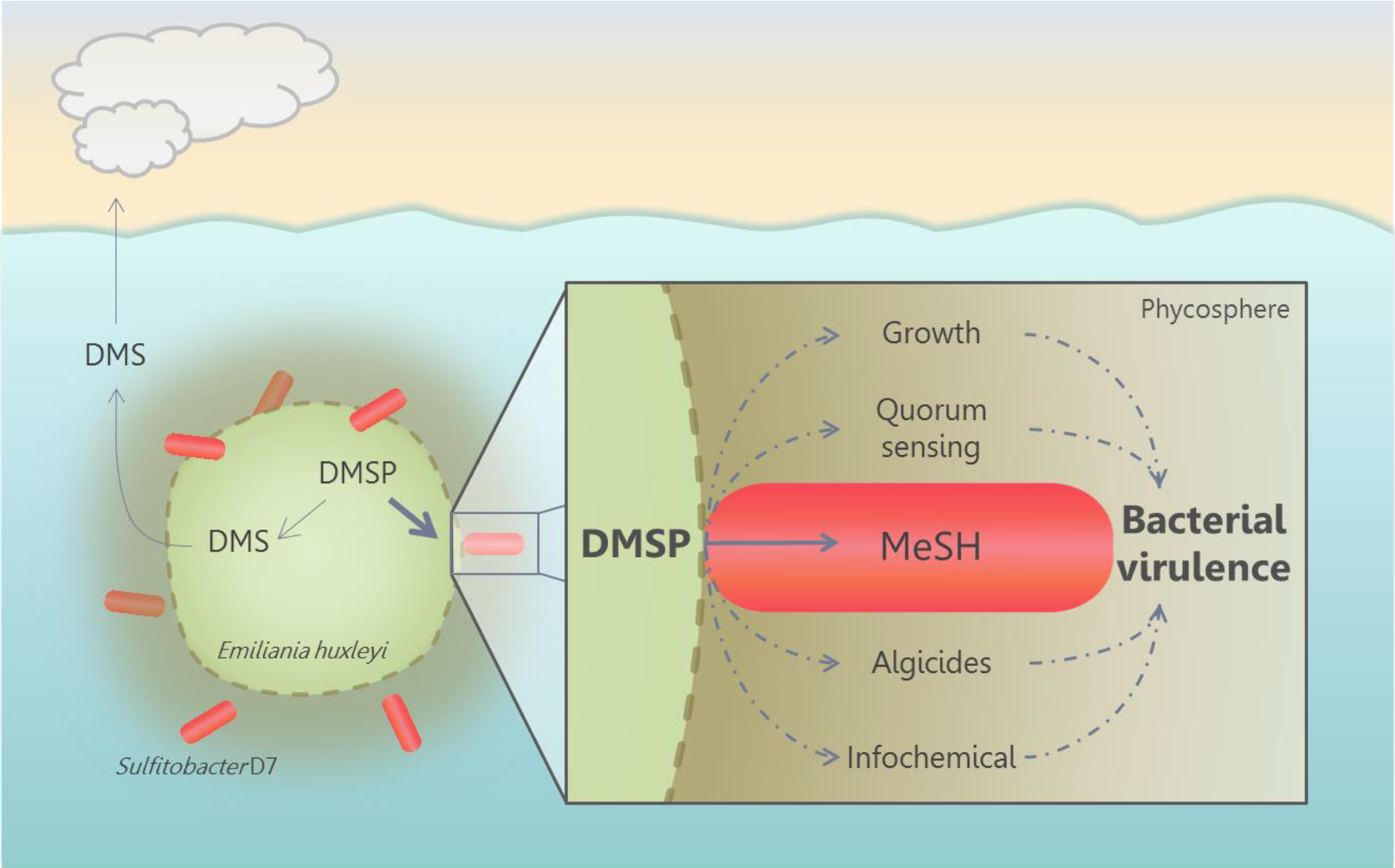
Conceptual model of the possible routes in which algal DMSP promotes bacterial virulence in *E. huxleyi* phycosphere. During interaction, *Sulfitobacter* D7 consumes *E. huxleyi*-derived DMSP and transforms it into MeSH, which facilitates bacterial growth. DMSP and its metabolic products can promote production of QS molecules (22) and bacterial algicides (52), which were proposed to be involved in bacterial virulence. Furthermore, DMSP may facilitate bacterial chemoattraction to algal cells (25,72). The algicidal effect of *Sulfitobacter* and other members of the *Roseobacter* clade (e.g. *P. inhibens* (15)) may have broader scale impact on the dynamics of *E. huxleyi* blooms. These blooms are an important source for DMSP and its cleavage product DMS, which is emitted to the atmosphere. By consuming large amounts of DMSP, bacteria may reduce DMS production by the algal DMSP-lyase (Alma1). Accordingly, we propose that the balance between competing DMSP catabolic pathways, driven by microbial interactions, may regulate oceanic sulfur cycling and feedback to the atmosphere.

## Materials and Methods

### Oceanographic cruise sampling and isolation of CAM bacterial consortium

Waters were collected from 61.5–61.87°N/33.5–34.1°W in June-July 2012, during the North Atlantic Virus Infection of Coccolithophore Expedition (NA-VICE; KN207–03), aboard the *R/V* Knorr (http://www.bco-dmo.org/project/2136). Samples were obtained from 5–6 depths using a Sea-Bird SBE 911plus CTD carrying 10 L Niskin bottles. Biomass from 1 to 2 L of seawater was pre-filtered through a 200 μm mesh, collected on 0.8 μm polycarbonate filters (Millipore), flash-frozen in liquid nitrogen, and stored at −80°C until further processing. Copepods were collected from surface waters (0–5 m) using 100 μm mesh nets on the 29^th^ of June (57.7°N, 32.2°W) and 11^th^ of July (61.9°N, 33.7°W) as described in Frada *et al*. (53). Single copepods were thoroughly washed with clean artificial seawater and kept at 4°C. Between 2 weeks to 1 month later, single copepod individuals were homogenized with a sterile pestle and inoculated into 2 mL of various *E. huxleyi* strains growing exponentially (53). Lysis of *E. huxleyi* strain NCMA 379 was observed within 1 week. The supernatant of the culture lysate was passed through a 0.45 µm filter and re-inoculated into *E. huxleyi* 379, resulting in the collapse of the culture. Addition of penicillin and streptomycin (20 units mL^−1^ and 20 µg mL^−1^, respectively) abolished culture lysis indicating the presence of bacterial pathogens (Supplementary Figure S1a). A suspension (<0.45 µm) of the culture lysate (copepod-associated microbiome, CAM) was kept at 4°C for further analyses.

### Isolation of *Sulfitobacter* D7 and *Marinobacter* D6

*Sulfitobacter* D7 and *Marinobacter* D6 were isolated from a co-culture of *E. huxleyi* 379 with CAM at 7 days of growth. Bacterial populations in co-cultures were stained with the live nucleic-acid fluorescent marker SYTO13 (Molecular probes). Two distinct sub-populations were observed in CAM-treated cultures (Supplementary Figure S1b) and were sorted at room temperature based on green fluorescence intensity (530/30 nm) in purity mode using a BD FACSAria™ II cell sorter equipped with a 488nm laser. Sorted populations were independently plated on marine agar plates (Difco) and incubated in the dark at 18°C. *Sulfitobacter* D7 and *Marinobacter* D6 were each isolated from a single colony and streaked three times from a single colony to ensure isolation of a single bacterial strain. For identification, DNA was extracted from a single colony of *Sulfitobacter* D7 using REDExtract-N-Amp Plant PCR kit (Sigma-Aldrich) according to the manufacturer’s instructions, and was used as a template for PCR with general primers for bacterial 16S rRNA: F- 5’-agtttgatcctggctcag-3’ and R- 5’-taccttgttacgacttcacccca-3’ (79). Amplicons were paired-end sequenced using the ABI 3730 DNA Analyzer and manually assembled. *Sulfitobacter* D7 and *Marinobacter* D6 were grown in marine broth (Difco) and stored in 15% glycerol at –80°C.

### Phylogenetic analysis

A multiple sequence alignment was generated using MUSCLE (80) with the default parameters. A maximum likelihood phylogeny was inferred using RAxML (81) under the GTRCAT model. Nodal support was estimated from a rapid bootstrap analysis with 1,000 replicates.

### *Sulfitobacter* D7 whole-genome sequencing and assembly

*Sulfitobacter* D7 genomic DNA was extracted using a DNeasy Blood & Tissue Kit (Qiagen) according to the manufacturer’s instructions. Genomic DNA was prepared for sequencing using the Nextera XT kit (Illumina, San Diego, CA) according to the manufacturer’s instructions. After processing, libraries were assessed for size using an Agilent TapeStation 2000 automated electrophoresis device (Agilent Technologies, Santa Clara, CA), and for concentration by a Qubit flurometer (Thermo Fisher Scientific Inc., Waltham, MA). Libraries were pooled in equimolar ratio and sequenced using an Illumina NextSeq500 sequencer, with paired-end 2x150 base reads. Library preparation and sequencing were performed at the DNA Services facility, University of Illinois at Chicago. Standard Pacific Biosciences large insert library preparation was performed. DNA was fragmented to approximately 20kb using Covaris G tubes. Fragmented DNA was enzymatically repaired and ligated to a PacBio adapter to form the SMRTbell Template. Templates larger than 10kb were BluePippin (Sage Science) size selected, depending on library yield and size. Templates were annealed to sequencing primer, bound to polymerase, and then bound to PacBio MagBeads and SMRTcell sequenced. Sequencing was performed at the Great Lakes Genomics Center at the University of Wisconsin – Milwaukee. *De novo* assembly was performed using the Spades assembler (82) on both raw Illumina and PacBio reads, with multiple k-mers specified as “-k 31,51,71,91”. Coverage levels were assessed by mapping raw Illumina reads back to the contigs with bowtie2 (83) and computing the coverage as the number of reads aligning per contig times the length of each read divided by the length of the contig. We assessed the relationship between coverage and cumulative assembly length over coverage-sorted contigs, and took 33% of the coverage level at half the total assembly length as a coverage threshold. Contigs with coverage less than this value or with a length shorter than 500 bp were removed. The sequence of *Sulfitobacter* D7 has been deposited in GenBank (BioProject PRJNA378866).

### Culture maintenance, axenization and bacterial infection

*E. huxleyi* strains were purchased from the National Center for Marine Algae (NCMA) and from the Roscoff culture collection (RCC) and maintained in filtered sea water (FSW). NCMA379, RCC1216, NCMA373 were cultured in f/2 medium (-Si) (84) and NCMA2090 was cultured in k/2 medium (-Tris, –Si) (85). Cultures were incubated at 18°C with a 16:8 h, light:dark illumination cycle. A light intensity of 100 µmol photons m^−2^ s^−1^ was provided by cool white LED lights. Cultures were made axenic by the following treatment (partially based on von Dassow *et al*. (86)): cells were gently washed with autoclaved FSW on sterile 1.2 µm nitrocellulose membrane filters (Millipore). Cells were transferred to algal growth media containing the following antibiotic mix: 20 µg mL^−1^ chloramphenicol, 120 units mL-^1^ polymyxin B, 40 units mL^−1^ penicillin and 40 µg mL^−1^ streptomycin. After 7 days the cultures were diluted into fresh algal growth media and the antibiotics mix was replenished. After another 7 days the cultures were diluted again into fresh algal growth media without antibiotics. For strains 1216, 373 and 2090, cultures were treated again with the following antibiotics mix: 50 µg mL^−1^ ampicillin, 25 µg mL^−1^ streptomycin and 5 µg mL^−1^ chloramphenicol. Cultures were transferred 1–2 times a week. After 2 weeks the cultures recovered and no bacteria could be detected by flow cytometry (see full description in the following section) or by plating on marine agar plates. Cultures were maintained with antibiotics and were transferred every 7–10 days. Prior to infection, *E. huxleyi* cultures were transferred 3–4 times to antibiotic-free algal growth media. For all experiments *E. huxleyi* cultures were infected at early exponential growth phase (2–4·10^5^ cell mL^−1^). For CAM infection, algal cultures were inoculated with 10^4^ bacteria mL^−1^. For *Sulfitobacter* D7 infection, bacteria were inoculated from a glycerol stock (kept at –80°C) into 1/2YTSS (2 gr yeast extract, 1.25 gr tryptone and 20 gr sea salts (Sigma-Aldrich) dissolved in 1 L DDW) and grown over-night at 28°C, 150 rpm. Bacteria were washed three times in FSW by centrifugation (10,000 g, 1 min). Algal cultures were inoculated at t = 0d with 10^3^ bacteria mL^−1^. In the experiment presented in Figure 5, *E. huxleyi* 2090 cultures were inoculated with 10^6^ bacteria mL^−1^. When noted, DMSP, glycerol or propionate were added at t = 0d. DMSP was synthesized according to Steinke *et al*. (32).

### Enumeration of algae and bacteria abundances and algal cell death by flow cytometry

Flow cytometry analyses were performed on Eclipse iCyt flowcytometer (Sony Biotechnology Inc., Champaign, IL, USA) equipped with 405 and 488 nm solid-state air-cooled lasers, and with standard optic filter set-up. *E. huxleyi* cells were identified by plotting the chlorophyll fluorescence (663–737 nm) against side scatter and were quantified by counting the high-chlorophyll events. For bacterial counts, samples were fixed with a final concentration of 0.5% glutaraldehyde for at least 30 min at 4°C, then plunged into liquid nitrogen and stored at −80°C until analysis. After thawing, samples were stained with SYBR gold (Invitrogen) that was diluted 1:10,000 in Tris–EDTA buffer, incubated for 20 min at 80°C and cooled to room temperature (RT). Samples were analyzed by flow cytometry (ex: 488 nm; em: 500–550 nm). For algal cell death analysis, samples were stained with a final concentration of 1 μM SYTOX Green (Invitrogen), incubated in the dark for 30 min at RT and analyzed by flow cytometry (ex: 488 nm; em: 500–550 nm). An unstained sample was used as control to eliminate the background signal.

### Scanning electron microscopy (SEM)

Samples of 0.5 mL were mixed with 0.5 mL of fixation medium (3% paraformaldehyde, 2% glutaraldehyde and 400mM NaCl, final) and stored at 4°C. Samples were adhered to silicon chips coated with poly-L-lysine (0.01%, Sigma-Aldrich). After three washes in 0.1M cacodylate buffer, samples were post-fixed with 1% OsO_4_ for 1 hr followed by three washes in 0.1M cacodylate buffer and three washes in milliQ water. Samples were dehydrated by a series of increasing concentration of ethanol (30% to 100%). Ethanol was replaced by liquid CO_2_ and critical point dried in BAL-TEC CPD 030. Lastly, samples were coated with gold/palladium (Edwards, S150) and imaged using High Tension Mode of XL30 ESEM.

### Enumeration of *E. huxleyi* and *Sulfitobacter* D7 by quantitative PCR (qPCR)

For environmental samples, genomic DNA was extracted using an adapted phenol– chloroform method previously described by Schroeder *et al*. (74). Filters were cut into small, easily dissolved pieces and placed in a 2 mL Eppendorf tube. Following addition of 800 μL of GTE buffer (50 mM glucose, 25 mM Tris-HCl (pH 8.0), and 10 mM EDTA), 10 μg mL^−1^ proteinase K, and 100 μL of 0.5 M filter-sterilized EDTA, samples were incubated at 65°C for 1–2 h. Following incubation, 200 μL of a 10% (vol/vol) stock solution of SDS was added and DNA was then purified by phenol extraction and ethanol precipitation. For lab samples, DNA was extracted using REDExtract-N-Amp Plant PCR kit (Sigma-Aldrich) according to the manufacturer’s instructions. *E. huxleyi* abundance was determined by qPCR for the cytochrome c oxidase subunit 3 (*cox3*) gene: Cox3F1: 5′-agctagaagccctttgaggtt-3’, Cox3R1: 5′-tccgaaatgatgacgagttgt-3’. *Sulfitobacter* D7 abundance was determined by qPCR for the 16S rRNA gene using primers designed in this study: 16S-D7bF: 5’-cttcggtggcgcagtgac-3’, 16S-D7bR: 5’-tcatccacaccttcctcccg-3’. The specificity of 16S-D7b primers were evaluated using TestPrime (https://www.arbsilva.de/search/testprime/) against the Silva SSU ref database (87). The primers matched only few *Sulfitobacter* sp. other than *Sulfitobacter* D7. All reactions were carried out in technical triplicates. For all reactions, Platinum SYBER Green qPCR SuperMix-UDG with ROX (Invitrogen) was used as described by the manufacturer. Reactions were performed on StepOnePlus™ real-time PCR Systems (Applied Biosystems) as follows: 50°C for 2 min, 95°C for 2 min, 40 cycles of 95°C for 15 s, 60°C for 30 s, followed by a melting curve analysis. Results were calibrated against serial dilutions of *E. huxleyi* (NCMA374 or NCMA2090) and *Sulfitobacter* D7 DNA at known concentrations, enabling exact enumeration of cell abundance. Samples showing multiple peaks in melting curve analysis or peaks that were not corresponding to the standard curves were discarded.

### Headspace analysis using solid-phase microextraction (SPME) coupled to gas chromatography mass spectrometry (GC-MS)

Headspaces of control, CAM- and *Sulfitobacter* D7-infected *E. huxleyi* 379 cultures after 10 days of growth were sampled for 15 min using an SPME DVB/Carboxen/PDMS fiber (Supelco, Bellefonte, PA, USA). Samples were manually stirred before absorption. For desorption, the fiber was kept in the injection port for 5 min at 260°C. Agilent 7090A gas chromatograph combined with a time-of-fight (TOF) Pegasus IV mass spectrometer (Leco, USA) was used for GC-MS analysis. Carrier gas (helium) was set as constant flow of 1.2 mL min^−1^. Chromatography was performed on an Rtx-5Sil MS column (30 m, 0.25 mm, i.d. 0.25 um) (Restek, Bellefonte, PA, USA). The GC oven temperature program was 45°C for 0.5 min followed by a 25°C min^−1^ ramp to a final temperature of 270°C with 3 min hold time. The temperatures of the transfer line and source were 250°C and 220°C, respectively. After a delay of 10 s, mass spectra were acquired at 20 scans s^−1^ with a mass range from 45 to 450 *m/z*. Peak detection and mass spectrum deconvolution were performed with ChromaTOF software (Leco). Identification was performed according to NIST library. Identification of DMDS was proofed by injection of commercial standard (Sigma-Aldrich).

### Evaluation of volatile organic sulfur compounds (VOSCs)

Samples were collected by small-volume gravity drip filtration (SVDF) (88) (see full description in the following section) and quickly diluted (1:10 in DDW) in a gas-tight vial. DMS, MeSH and DMDS levels were determined using Eclipse 4660 Purge-and-Trap Sample Concentrator system equipped with Autosampler (OI Analytical). Separation and detection were done using gas chromatography-flame photometric detector (GC-FPD, HP 5890) equipped with RT-XL sulfur column (Restek). The GC oven temperature program was 100°C for 1 min followed by a 70°C min^−1^ ramp to a final temperature of 240°C with 7 min hold time. All measurements were compared to standards (Sigma-Aldrich) (Supplementary Figure S6). For calibration curves, DMS and DMDS were diluted in DDW to known concentrations. For MeSH standard we used MeS^−^Na^+^ dissolved in DDW and added HCl in 1:1 ratio by injection through the septa of the vials. We could not quantify MeSH since part of it was oxidized to DMDS during the procedure (Supplementary Figure S6c). Therefore, MeSH abundance is presented as square-root of the area of the peak corresponding to MeSH. No VOSCs were present in blank (DDW) samples.

### Determination of DMSP concentration

#### Lab experiments

Samples for dissolved DMSP (DMSP_d_) were obtained by small-volume gravity drip filtration (SVDF) (88). *E. huxleyi* cultures were filtered through Whatman GF/F filters by gravity using filtration towers. Filtrates (∼3 mL) were acidified to 1.5% HCl for DMSP_d_ preservation and stored at 4°C for > 24 h. Samples were diluted (typically 1:100) in DDW and DMSP_d_ was hydrolyzed to DMS by adding NaOH in a final concentration of 0.45M and incubated for 1 h at room temperature in the dark. Glycine buffer (pH = 3) was added to a final concentration of 0.8 M for neutralization (pH = 8–9, final). Samples were measured for DMS.

#### Cruise

Collection of water samples is described in the first section of the Materials and Methods. For determining DMSP_d_, ≤ 20 mL were collected by SVDF and the filtrate was acidified with 50% sulfuric acid (10 μL per 1 mL of sample). Sample preparation was conducted at room temperature. All DMSP samples were stored at 4°C until analysis. Upon analysis, the samples were base-hydrolyzed in strong alkali (sodium hydroxide; final concentration 2 mol L^−1^) and analyzed for DMS. Instrumental determination of DMSP (as DMS) was carried out using Membrane Inlet Mass Spectrometry (MIMS) (89,90) system that comprised of a Pfeiffer Vacuum quadrupole mass spectrometer equipped with a HiCube 80 pumping station, QMA 200 analyzer and flow-through silicone capillary membrane inlet (Bay Instruments, Easton, Maryland). The inlet consisted of a glass vacuum line incorporating a U-tube and support for the 0.51 mm ID Silastic tubing membrane and 0.5 mm ID stainless steel capillary supply lines. The sample was pumped through the inlet system at 1.5 mL min^−1^ using a Gilson Minipuls 3 peristaltic pump. Prior to entering the membrane, the sample passed through a 75 cm length of capillary tubing immersed in a thermostated water bath (VWR, Suwanee, GA) and held at 30°C to ensure constant temperature (and membrane permeability) as the sample passed through the membrane. The U-tube section of the vacuum line (located between the membrane inlet and mass spectrometer) was immersed in an isopropanol bath (held at < –45°C) to remove water vapor from the gas stream prior to introduction of the stream into the mass spectrometer. In this configuration, the system maintained an operating vacuum pressure of 2.0 (± 0.2) ·10^−5^ mbar. The sample liquid was pumped from the bottom of the sample test tube and through the membrane until the mass spectrometer signal stabilized (typically a minimum of six minutes). DMS was monitored semi-continuously by scanning at *m/z* 62 for 5 seconds every 15 seconds using the secondary electron multiplier detector. Calibration of the MIMS instrument was carried out with freshly-prepared base-hydrolyzed DMSP standards made using ESAW synthetic seawater and commercially-available DMSP powder (Research Plus, Bayonne, New Jersey). The detection limit for the system was 0.2 nmol L^−1^.

### *Sulfitobacter* D7 growth in conditioned media (CM) and minimal medium (MM) supplemented with DMSP

*Sulfitobacter* D7 were grown over-night in 1/2YTSS at 28°C. Bacteria were washed three times in artificial sea water (ASW) (91) by centrifugation (10,000 g, 1 min). Media were inoculated with 10^4^ bacteria mL^−1^. CM were obtained from mono-cultures of *E. huxleyi* strains by SVDF (88). This method was chosen in order to prevent lysis of algal cells during the procedure and release of intracellular components. Following SVDF, media were filtered through 0.22 µm using syringe filters. In the experiment presented in Figure 4b bacterial growth was followed for 24h. MM was based on ASW supplemented with basal medium (-Tris) (BM, containing essential nutrients) (92) and vitamin mix (93). In the experiment presented in Supplementary Table S2 the MM was supplemented with 1 gr L-^1^ glycerol and 70 µM DMSP (synthesized according to Steinke *et al*. (32)). Bacterial growth, DMSP_d_ and VOSCs levels were measured at t = 0h and t = 24h. In the experiment presented in Supplementary Figure S9 the MM was supplemented with 0.01 gr L^−1^ glycerol, 0.5 mM NaNO_3_, metal mix of k/2 medium (85) and different concentrations of DMSP. Bacterial growth was measured at t = 16h.

### Statistical analyses

For all time-course experiments, significant differences in the various parameters were determined using a one-way/two-way repeated measures ANOVA. In other experiments, differences were tested by a one-way ANOVA. Tukey post-hoc tests were used when more than two levels of a factor were compared.

## Acknowledgments

We thank the chief scientist of the NA-VICE cruise, Kay D. Bidle (Rutgers University), and the captain and crew of the *R/V* Knorr, the Marine Facilities and Operations at the Woods Hole Oceanographic Institution for assistance and cooperation at sea. We thank Uria Alcolombri (ETH, Zurich) and Alon Amrani (The Hebrew University of Jerusalem, Israel) for their assistance in GC analyses. We thank Shifra Ben-Dor for her contribution to the phylogenetic analysis. We thank Guy Schleyer for his great assistance and constructive feedback on the manuscript. We thank Assaf R. Gavish for fruitful discussions and assistance in graphics. We thank the Rieger Foundation for granting a JNF Fellowship in Environmental Studies to NBG. All SEM studies were conducted at the Electron Microscopy Unit of the Weizmann Institute of Science, supported in part by the Irving and Cherna Moskowitz Center for Nano and Bio-Nano Imaging. This research was supported by the European Research Council (ERC) StG (INFOTROPHIC grant # 280991), CoG (VIROCELLSPHERE grant # 681715) and NSF grants OCE-1061883 (to AV), OCE-1061876 (to GRD), OCE-1436458 and OCE-1428915 (to PAL).

